# Past, present and future in the geographical distribution of Mexican tepezmaite cycads: genus *Ceratozamia*

**DOI:** 10.1101/2023.03.23.533928

**Authors:** Jorge Antonio Gómez-Díaz, César Isidro Carvajal-Hernández, Wesley Dáttilo

## Abstract

*Ceratozamia morettii*, *C. brevifrons* and *C. tenuis* are cycads considered endangered in montane forests in the center of Veracruz state. However, the amount of theoretical and empirical information available on the historical distribution of these species and how they could be affected in the future by the effects of climate change is still limited. Our objective was to generate information on the spatial distribution of the species since the last glacial maximum, present and future. To map the spatial distribution of species, we created a potential distribution model for each species. The spatial data used for the models included 19 bioclimatic data variables in the present, at the last glacial maximum using two models (CCSM4 and MIROC), and in the future (2080) using two models of the RCP 8.5 scenario of climate change (HadGEM2-CC and MIROC5). We found that each species occupies a unique ecoregion and climatic niche. *Ceratozamia morettii* and *C*. *tenuis* have a similar pattern with an expansion of their distribution area since the last glacial maximum with the larger distribution area in the present, and with a projected reduction in their distribution under future climatic conditions. For *C. brevifrons,* we also showed an increase in their distributional area since the last glacial maximum and we showed that this expansion will continue under future climatic conditions when the species will reach its maximum distributional area. Projections about the future of these endemic cycad species show changes in their habitat highlighting that temperate zone species will face imminent extinction if no effort is made to protect them. On the other hand, the tropical climate species will apparently be favored.

## Introduction

Cycads are an endangered plant group that is distributed in tropical and subtropical regions of the world [1]. They are considered living fossils and they evolved from the ancient “seed ferns” of the late Paleozoic. According to [2], there are 370 described species of cycads in the world. In Mexico, 54 species of cycads are recognized in three genera: *Ceratozamia*, *Dioon* and *Zamia* [3], however new species have recently been described. Mexico is second place in the diversity of cycads after Australia with 80% endemic species of the country [4].

Cycads live in tropical and subtropical environments, from humid jungles, dry jungles, cloud forests, pine-oak forests, and scrublands [4, 5]. Unlike other non-flowering plants, cycads are pollinated by primitive insects (Coleoptera and Thysanoptera) that some have their origin in the mid-Cretaceous [6, 7]. The seeds of cycads have been used as food supplementing the diet of people in times of scarcity, but in Mexico, its main use is as ornamental, the leaves of some cycads are used to decorate churches during religious festivals [8]. Many species of cycads are removed from their natural habitats by collectors and plant traders [4], added to the reduction of their habitats, has caused that currently more than 60% cycad species in the world are threatened [5].

Among the cycads, there is the *Ceratozamia* genus (Zamiaceae) which is exclusively distributed in the nootropics [9,10]. Most of the species of *Ceratozamia* are in the mountainous regions of Mexico, specifically on the mountain range of the Sierra Madre Oriental in which are the highest diversity of this genus [9,10].

Within the genus *Ceratozamia* is *C. brevifrons* (Fig. 1a), *C. morettii* (Fig. 1b) and *C. tenuis* (Fig. 1c), three micro endemic species from central Veracruz, known by the local name “Tepezmaite”, which are at risk of disappearance [8]. All three species are considered endangered by International Union for Conservation of Nature (IUCN) and are listed in section I of CITES [11,12]. On the other hand, *C. morettii* and *C. brevifrons* (the latter as synonym are *C. mexicana*) are also protected by Mexican laws [13, 14]. *C. tenuis* has not been evaluated by Mexican laws due to its recent lectotype assignment in 2016 [15], previously considered a synonym of the species *C. mexicana*.

**Fig. 1.**
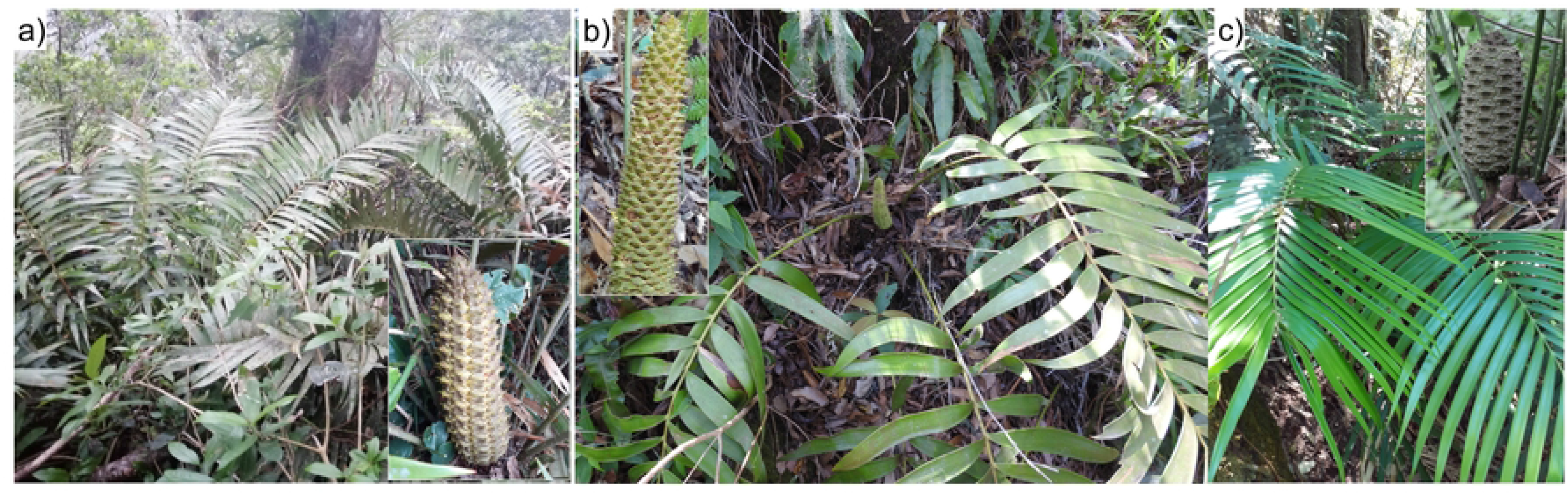
Tepezmaite cycads. a) *Ceratozamia brevifrons* with female cone; b) *C*. *morettii* with male cone; c) *C*. *tenuis* with female cone)

The three species grow on humid mountain forests and oak, which are the most threatened vegetation types in Mexico [16]. Besides, the habitat has strong pressures caused by agricultural and livestock activities [17]. Therefore, these species are of particular interest for conservation strategies as well as management purposes.

In general, IUCN range polygons are frequently used to make decisions for the conservation of species [18,19]. However, this methodology does not consider the effects of environmental variables that could influence the species and sometimes can severely underestimate the real-world occupancy [20–22]. Therefore, it has been recognized that the IUCN range polygons are imprecise and overestimate the size of the distribution of the species, with the consequences for their conservation that this methodology entails [23]. Species distribution modelling has been used as a tool to determine the suitability of the ecological niche of the species (i.e., species habitat requirements necessary for long-term population persistence). This is because niche ecological models were developed to predict species distributions from presences [24], that together with the ease and progress that has been made in this area has led to their greater use.

In addition to the geographic information provided by species distribution modelling, it has also been used for the conservation of different taxonomic groups such as (birds, mammals, and plants). However, this tool has been used little for cycads, since only some works address these species from the perspective of their conservation [25]. The evaluation of the species concerning the past conditions in the glaciations, allow us to understand how the contractions in the distribution of the species occurred under adverse weather conditions [26]. In this sense, in old groups as the cycads, which come from more than 200 ma [27], the information generated by species distribution modelling can help us to understand know how they developed in conditions different from the current ones, in addition to the fact that currently, most species are in danger of extinction [5].

In general, the species that inhabit mountainous areas, such as the tepezmaite’s cycads, are the most vulnerable to the effects of climate change [28,29]. The main effect of climate change on these species is found in the movement of their populations since they seek their preferred climatic niches over time, which is why montane populations migrate to new higher areas avoiding changes in the climate [29,30]. However, the high rates of deforestation and transformation of montane habitats wreak havoc on the survival of populations at ever-increasing rates, if climate change is added to this, the result is populations that are increasingly fragmented and at greater risk of extinction [29,31]. Therefore, one of the goals of the species distribution models is to prevent and forecast changes in the distribution patterns of threatened species in the face of the threat of climate change, with which the necessary actions could be taken to help in their conservation [29, 32].

Unfortunately, it is not known what the effects of climate change could be on the distribution of tepezmaite cycads, which is one more threat to the conservation of these species [28]. In fact, according to the IUCN, conducting assessments regarding the effect of climate change on species is important, to anticipate the effects that the future climate may have on current populations and thus anticipate and act effectively for their conservation [33].

Therefore, the objective of this study was to know the changes in the past distribution (last glacial maximum), present and future (under the effects of climate change) of the tepezmaite cycads (*C*. *brevifrons*, *C*. *morettii* and *C*. *tenuis*), to understand what is the history that the species have had and how they could be affected in the future by the effects of climate change. Specifically, we seek to know: i) which are the environmental variables that most influence the model of the ecological niche of each species, ii) which was the distribution of each species in the last glacial maximum, iii) which is the present distribution of each species and iv) what are the effects of climate change on species for the year 2080.

## Methods

### Study area

Our study area lays in the center of Veracruz state, Mexico in the Sierra de Chiconquiaco because of the micro endemic species (Fig. 2). Located in the Sierra Madre Oriental, the Sierra de Chiconquiaco has a complex system of micro- environments and great cultural wealth (Fig. 2). This is due to the short distance that exists between the coastal zone and the elevations of the mountain range, which rises from thirty-five to almost 3,000 m a.s.l. [34].

**Fig. 2.**
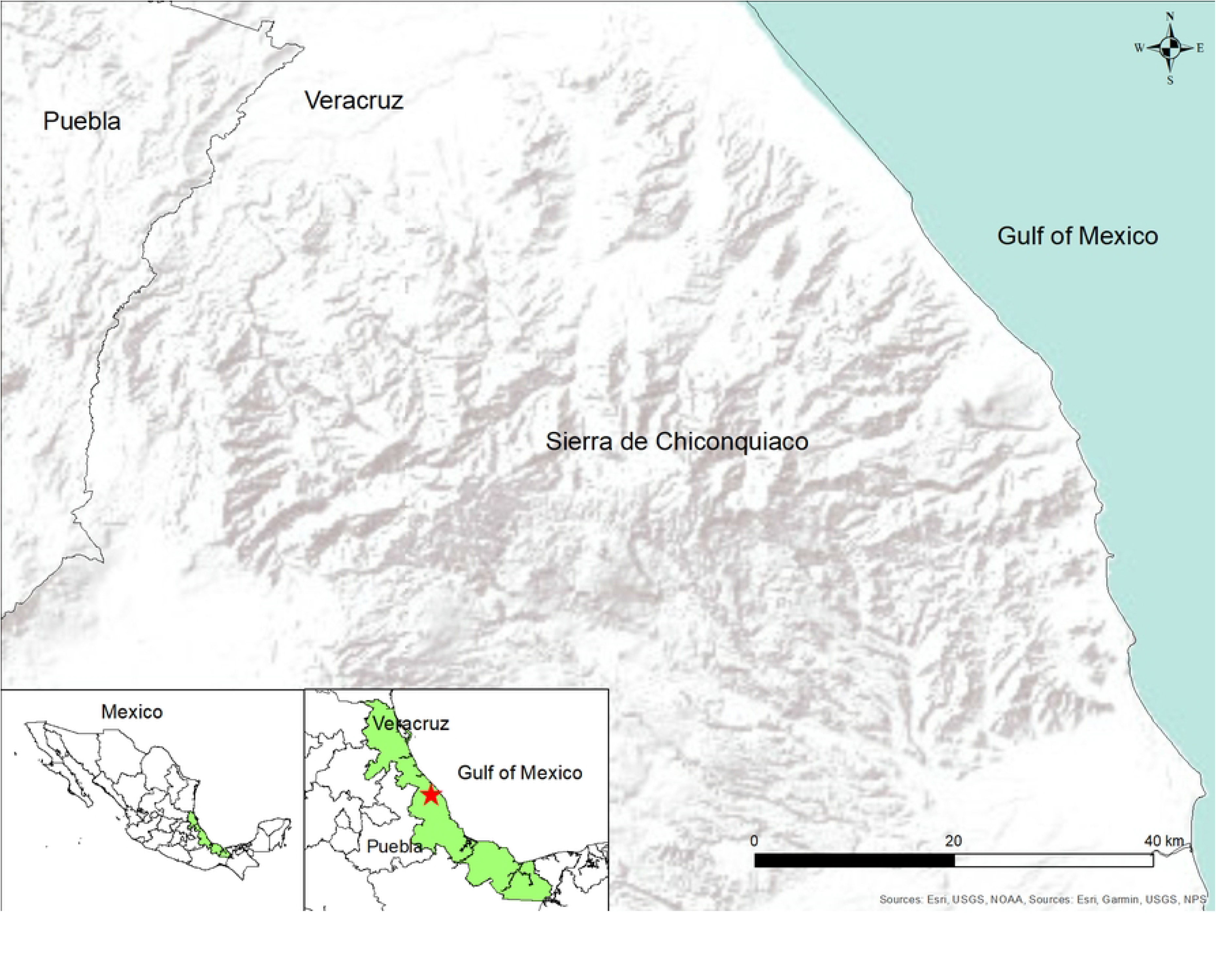
Study area in the Sierra de Chiconquiaco, Veracruz, Mexico

The biodiversity of this area is threatened due to several activities such as deforestation, forest fires, livestock and agriculture, particularly in mountains [35]. This area is topographically and climatically complex with seven types of vegetation and possesses a high plant diversity with 3019 species, including thirty-six endemic, 57 protected by Mexican laws and 195 listed in CITES (including six cycads) [33].

### Mapping the species

To map the spatial distribution of species, we obtained records of the studied *Ceratozamia* species from our data in the field, data from the XAL, XALU and MEXU herbaria, data from the literature and the Global Biodiversity Information Fund (GBIF), as well as corroboration of localities in the field. We found 14 occurrences for *C*. *brevifrons*, 16 for *C*. *morettii* and 26 for *C*. *tenuis*. The coordinates were cleaned up (i.e., doubtful records based on known distributional ranges) using the *clean_coordinates* function from the *CoordinateCleaner* R package [36]. The data was cleaned to have only one presence per km^2^. In the end, only thirteen records of *C*. *brevifrons* remained 16 of *C*. *morettii* and 21 of *C*. *tenuis*, which entered the distribution model. Only georeferenced records were chosen. Once the database with the points of presence was completed, a potential distribution model for each species was conducted.

The spatial data used for modelling included nineteen bioclimatic data variables from the CHELSA v.1.2 database at a resolution of 30 arc seconds [37]. The CHELSA bioclimatic variables have proven to be the most suitable in mountainous and tropical areas [38]. For the present, the last glacial maximum (∼ 22,000 years ago; past henceforth) and the scenario of future climate change (2080; future henceforth) projections, we consider the same bioclimatic variables.

In the case of the past, we used the median of the predictions of two Global Circulation Models (CGMs) models (CCSM4 and MIROC). In the case of future, we also used the median of two CGMs models (HadGEM2-CC and MIROC5) which are commonly used and recommended for Mexico based [39] on the RCP (Representative Concentration Pathway) 8.5 scenario of climate change of the IPCC (Intergovernmental Panel on Climate Change). We only considered the RCP 8.5 (936 ppm of CO2) scenario because represents a continuous increase in global emissions throughout the 21st century [40]. Therefore, if any of the species could survive in this scenario is highly likely to endure future climate outcomes.

We consider the same bioclimatic variables for the distribution modelling at the last glacial maximum, present and under climate change scenario. An area of exploration of the variables (M) was calibrated, using the terrestrial ecoregions of The Nature Conservancy that are based on those created by [41]. For each species, the ecoregions that had at least one presence record were selected. The variables were trimmed to the M of each species.

Also, the environmental variables were revised to avoid multiple collinearities between the variables using the variance inflation factor (VIF) analysis. Variables with a VIF value > 10 were eliminated using the exclude function from the R usdm version 1.1-18 package [42]. VIF indicates the degree to which standard errors are inflated due to levels of multicollinearity. VIF values > 10 were taken as indicative of problematic collinearity/redundancy [43] since high collinearity could lead to poor model performance and misinterpretations [38]. Of the 19 variables, only six variables did not present multicollinearity (VIF <10) for *C*. *brevifrons*, and five for both *C*. *morettii* and *C*. *tenuis* (Table 1).

**Table 1.** Bioclimatic variables (names and units) were used as predictors in the species distribution models of tepezmaite cycads (Ceratozamia brevifrons, C. morettii and C. tenuis).

| Variable | Name | Species |
| --- | --- | --- |
| Bio 2 | Mean Diurnal Range | *Ceratozamia brevifrons* |
| Bio 3 | Isothermality | *Ceratozamia morettii* |
| Bio 4 | Temperature Seasonality | *Ceratozamia brevifrons*, *C. morettii* |
| Bio 7 | Temperature Annual Range | *Ceratozamia tenuis* |
| Bio 8 | Mean Temperature of Wettest Quarter | *Ceratozamia morettii* |
| Bio 9 | Mean Temperature of Driest Quarter | *Ceratozamia brevifrons* |
| Bio 11 | Mean Temperature of Coldest Quarter | *Ceratozamia brevifrons*, *C. tenuis* |
| Bio 13 | Precipitation of Wettest Month | *Ceratozamia brevifrons*, *C. morettii*,<br>*C. tenuis* |
| Bio 14 | Precipitation of Driest Month | *Ceratozamia tenuis* |
| Bio 18 | Precipitation of Warmest Quarter | *Ceratozamia brevifrons*, *C. morettii*,<br>*C. tenuis* |

For each species, an Ensemble species distribution model (ESDM) was generated assembling all the methods available in the R package *SSDM* using the function *ensemble_modelling*. The methods included: Generalized linear model (GLM), Generalized additive model (GAM), Multivariate adaptive regression splines (MARS), Generalized boosted regressions model (GBM), Classification tree analysis (CTA), Random forest (RF), Maximum entropy (MAXENT), Artificial neural network (ANN), and Support vector machines (SVM). To generate the binary map of each species, the sensitivity-specificity equality (SES) metric was chosen [44]. Subsequently, AUC (area under the curve) values were calculated to describe the model performance or predictive accuracy. In this study, all statistical analyzes were performed using the programming language R v.3.6.3 [45].

## Results

In general, all the models for the *Ceratozamia* species have an AUC above 0.9 which means that the prediction of each model was accurate (Table 2). Despite the three species living extremely near each other (∼10 km) and having an exceptionally low distribution area in central Veracruz (Fig. 3), each one occupies a unique ecoregion and climatic niche. In this case, the ESDM for each species shows that they have a difference in the importance of bioclimatic variables influencing the distribution of the *Ceratozamia*. In the case of *C. brevifrons*, the most important variables were Bio 4 (41%), Bio2 (17%) and Bio 13 (16%), for *C. morettii* the most important variables were Bio 8 (29%), Bio 13 (24%) and Bio 4 (23%), and for *C. tenuis* were Bio 14 (29%), Bio 11 (24%) and Bio 18 (20%; Table 3). The only variables shared for the three species are Bio 13 and Bio 18.

**Fig. 3.**
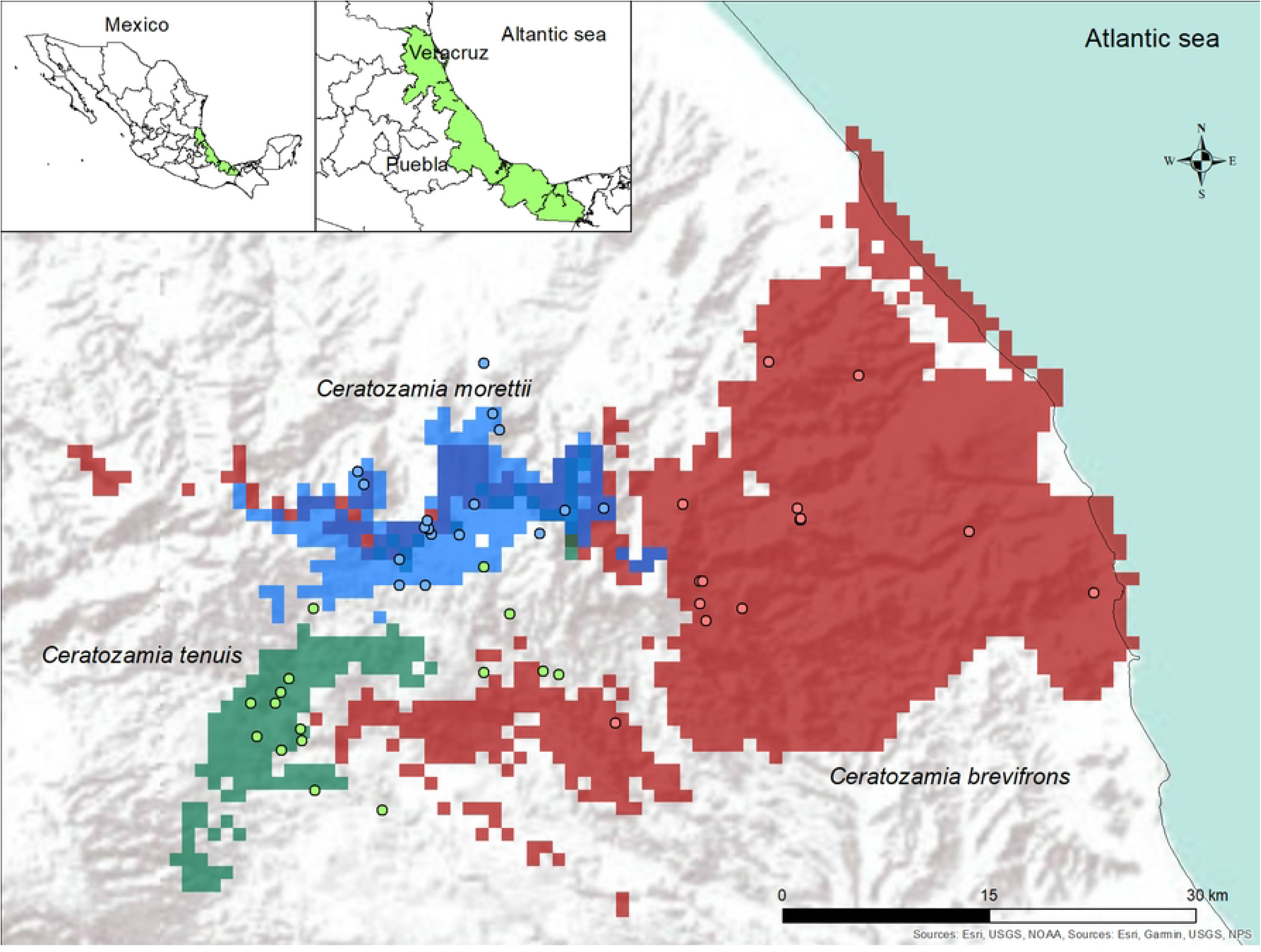
Potential distribution of the studied *Ceratozamia* species with the locations of occurrence of each species (points) in the Sierra de Chiconquiaco, Veracruz, Mexico, the grey shadow represents topographic relief

**Table 2.**
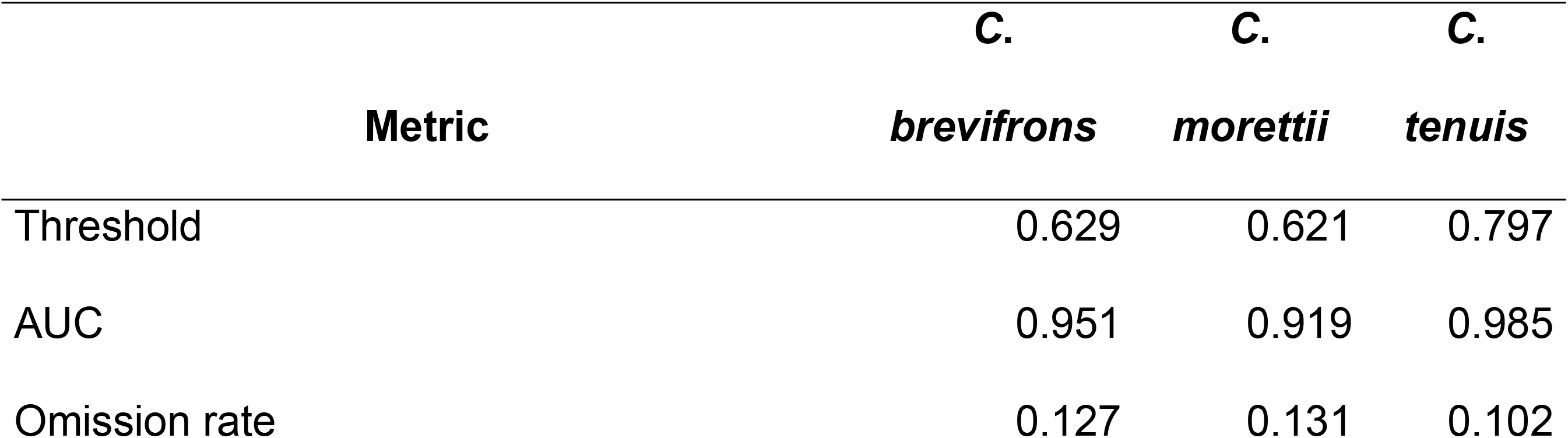
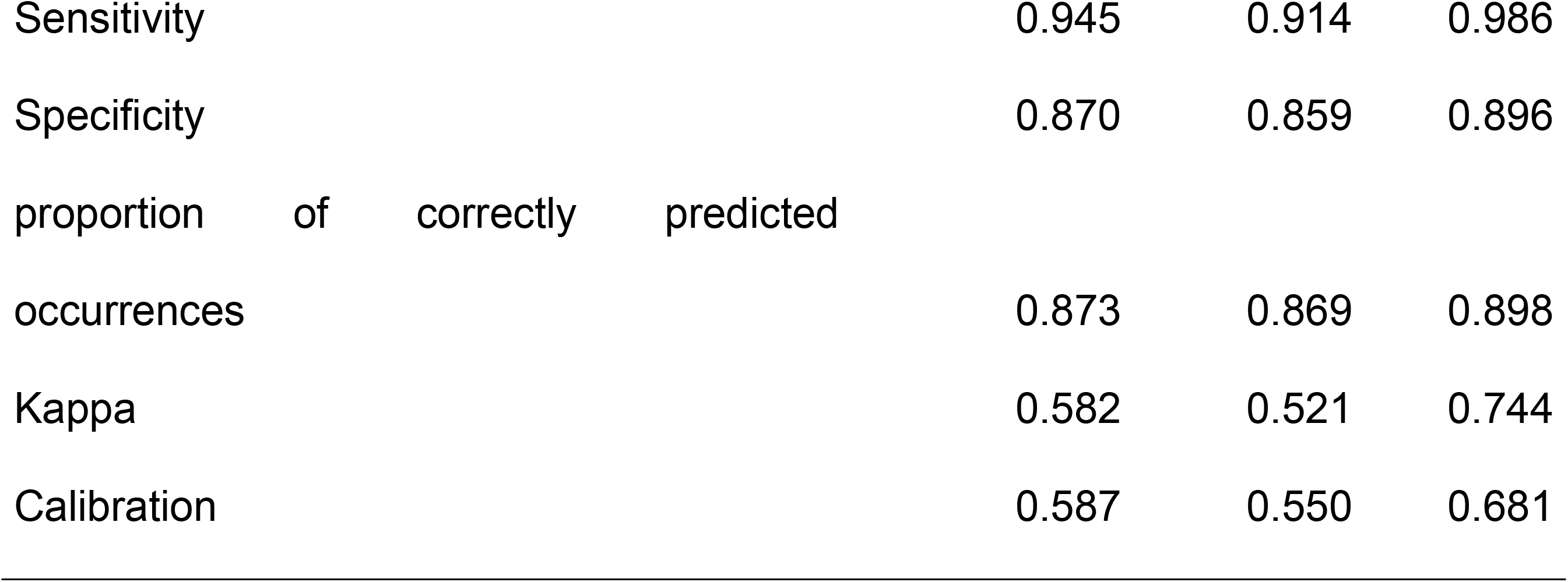
The threshold to categorize the suitability and evaluation metrics for the ESDM for each Ceratozamia studied.

**Table 3.**
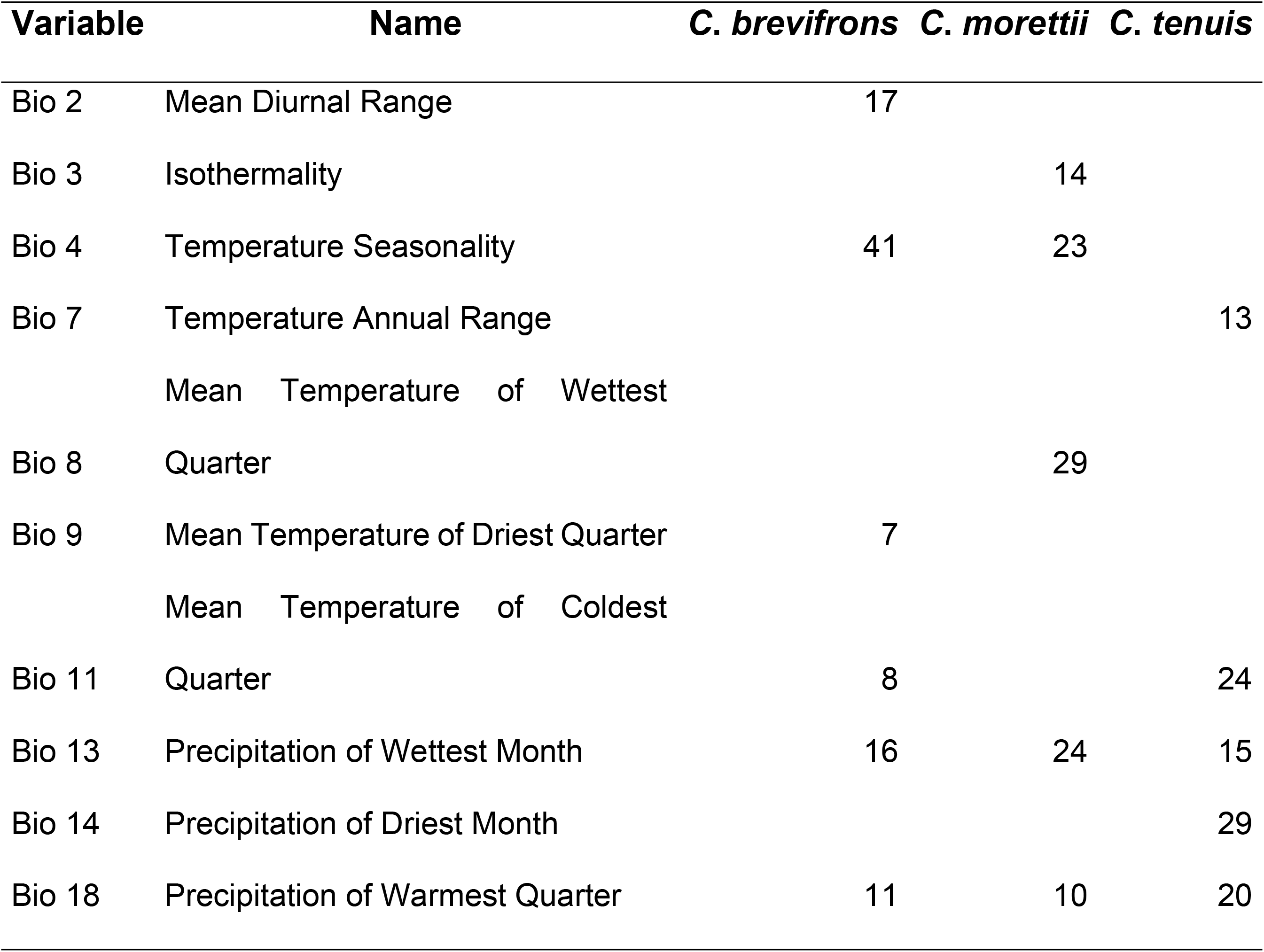
Variable importance (%) for the ensemble species distribution model of each of the *Ceratozamia* studied.

### Ceratozamia brevifrons

Modelling the last glacial maximum (LGM) suggested a slight decrease (Table 4) of occurrence for *C. brevifrons* and the species was restricted to the mountains of the area (Fig. 4a). The species had an increase of 155% in its geographical distribution from past to present. The present potential distribution of *C. brevifrons* covered the area of central Veracruz, with the most favorable conditions on the east of the Sierra de Chiconquiaco in some mountains and near the coast (Table 4, Fig. 4b). Of the three species in this study, this is the one with the lowest elevation (500 m a.s.l.). Less favorable areas for *C. brevifrons* were defined in the forests at the north and south of the Sierra de Chiconquiaco. Though, under the future scenario, an increase of 60% in the geographical range of the species is expected (Table 4 and 5, Fig. 4c).

**Fig. 4.**
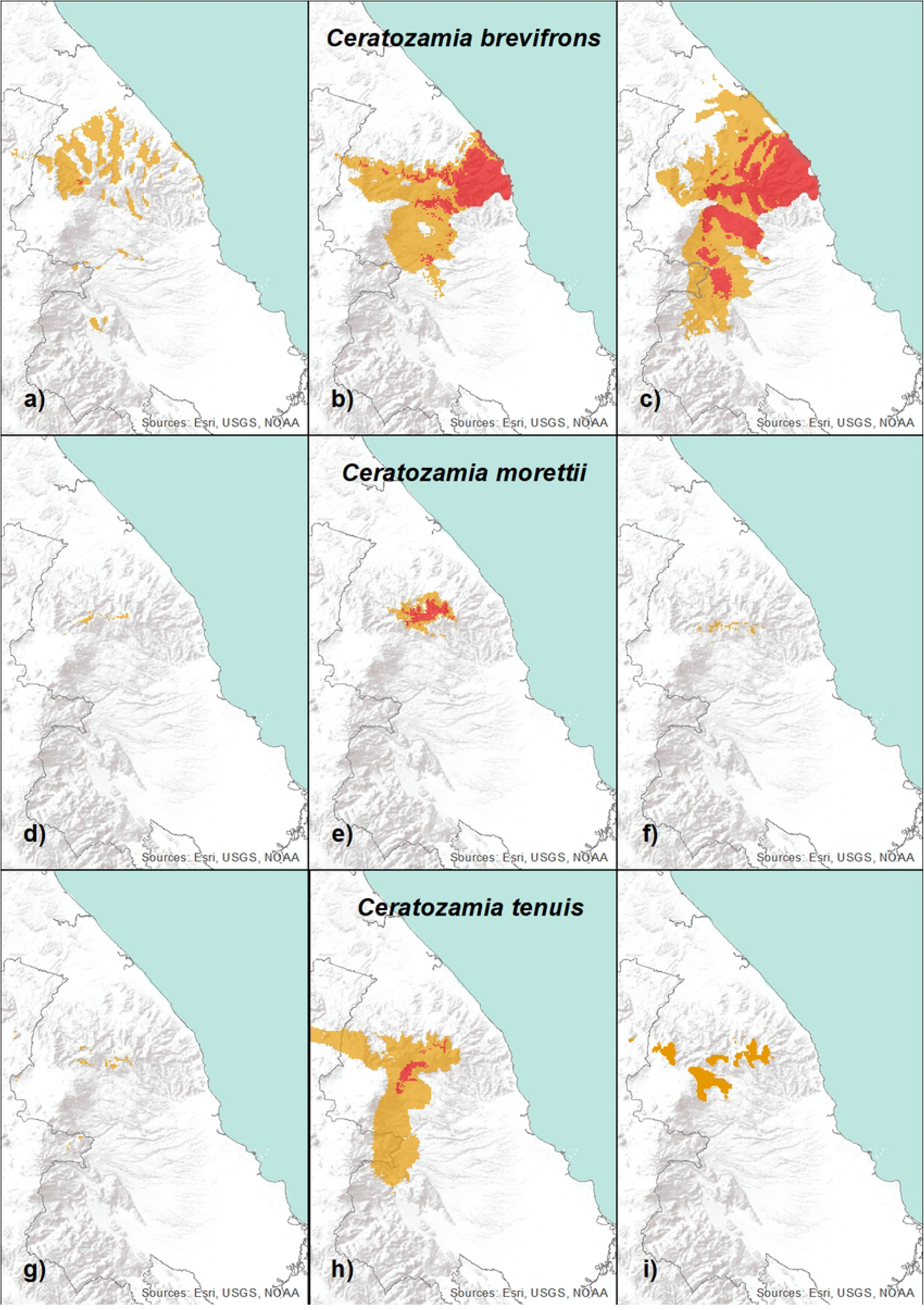
Potential distribution of *Ceratozamia brevifrons* a) in the past; b) present; c) future; *C. morettii* d) in the past; e) present; f) future; and *C. tenuis* g) in the past; h) present; i) future. Red: high suitability; Orange: medium

**Table 4.** Projected suitability area (km^2^) of the three species of Ceratozamia studied in the various times past (last maximum glacial), present, future (2080).

| Time | Suitability | *C. brevifrons* | *C. morettii* | *C. tenuis* |
| --- | --- | --- | --- | --- |
| Past | High | 5 | 0 | 0 |
|  | Medium | 1409 | 41 | 74 |
| Present | High | 1096 | 166 | 121 |
|  | Medium | 2515 | 279 | 2917 |
| Future | High | 2214 | 0 | 0 |
|  | Medium | 3581 | 54 | 529 |

### Ceratozamia morettii

Modelling the last glacial maximum (LGM) suggested a decrease of occurrence for *C. morettii* (Table 4, Fig. 4d). The species had an increase of 985% in its geographical range from the past to the present (Table 5). While the present potential distribution of *C. morettii* covered the area of central Veracruz, with the most favorable conditions in the Sierra de Chiconquiaco (Table 4, Fig. 4e). In this area, there are a large number of records of the species. The least favorable areas for *C. morettii* were found in the cloud forest at the south and west of the Sierra de Chiconquiaco. But, under the climate change scenario of the models HadGEM2-CC and MIROC 5, a reduction of 88% in the species’ range of distribution is expected (Table 4 and 5, Fig. 4f). Except for limited high elevated areas in the mountains of the Sierra de Chiconquiaco, the species practically disappears.

**Table 5.** Percentage of change between the different periods of the studied tepezmaite cycads.

| Change | *C. brevifrons* | *C. morettii* | *C. tenuis* |
| --- | --- | --- | --- |
| Past to present | 155% | 985% | 4005% |
| Present to future | 60% | -88% | -83% |
| Past to future | 310% | 32% | 615% |

### Ceratozamia tenuis

Modelling the last glacial maximum (LGM) suggested a decrease of occurrence for *C. tenuis* (Table 4, Fig. 4g). The species had an increase of 4005% in its geographical range from past to present (Table 5). On the other hand, the present potential distribution of *C. tenuis* covered the area of central Veracruz, with the most favorable conditions in the cloud forests in the south of the Sierra de Chiconquiaco (Table 4, Fig. 4h). In this area, there are a large number of records of the species. The least favorable areas for *C. tenuis* were found in the cloud forest at the south and west of the Sierra de Chiconquiaco. But, under the climate change scenario, a reduction of 83% of the species’ range of distribution is expected (Table 4 and 5, Fig. 4i). Except for limited areas in mountainous areas on the south of the Sierra de Chiconquiaco, the species practically disappears.

## Discussion

We estimated the past, present, and future distribution of three endemic cycads of Veracruz (*C*. *brevifrons*, *C*. *morettii* and *C*. *tenuis*). We provide a quantitative assessment of the Spatio-temporal changes in the potential distribution of these threatened species since the last glacial maximum and as a consequence of global climate change. In summary, we found: (a) a general increase distributional area of the species since the last glacial maximum, (b) a decrease in the range occupied by *C*. *morettii* and *C*. *tenuis* under future scenarios and (c) an increase in the range occupied by *C*. *brevifrons* in future scenarios.

We found that the *Ceratozamia* had dynamics in their range of distribution with expansions from the LGM. The modelled distributions showed a spatial separation among the three species since the LGM, which is a pattern found also for the conifer *Podocarpus* (Podocarpaceae) in Mexico [46]. This could be explained by the difference between the bioclimatic variables that affect each species. The models indicate that the precipitations of the wettest month (Bio 13) and the warmest quarter (Bio 18) were the only variables shared for the three species of *Ceratozamia*. Moreover, we found that the temperature seasonality (Bio 4) was very important for *C. brevifrons*, while the average temperature of the wettest quarter (Bio 8) was for *C. morettii* and precipitation of the driest month (Bio 14) was for *C. tenuis*. These findings are accordingly to [25] who found for *C*. *miqueliana*, a species for tropical rain forest from southern Veracruz, that precipitation in the driest month was also a variable that explains the distribution of the species.

According to the Pleistocene refuge hypothesis, in places where climatic changes were not so extreme, many species would find refuge and would be restricted to these areas [47]. This is consistent with those observed in this study since using the distribution models, we observed that there was an effect on the distribution of *Ceratozamia* species due to past climatic changes in the Quaternary (LGM). Specifically, we observed that montane species (*C*. *morettii* and *C*. *tenuis*) encountered cloud forest conditions in highly restricted areas of the Chiconquiaco mountain range and that by the humid forest model they moved to lowlands in the LGM [48]. A population movement towards lowlands from highlands was expected since that during the LGM these lowlands would serve as refuges with limited habitat suitability, all this under the Pleistocene refuge hypothesis which postulates climate-driven expansions and contractions of species ranges [48]. We found support for that hypothesis, observing that the populations of the three *Ceratozamia* species had limited habitat suitability during the LGM.

The above was more marked in the species *C. morettii* and *C. tenuis* under the hypothesis of long-term in situ persistence because the cloud forest currently has a fragmented distribution in our study region. According to this hypothesis, the populations of *Ceratozamia* would be likely geographically structured and persistent (multiple glacial refuges), with no signs of demographic expansion before IGM and limited gene flow [46]. Comparable results of local persistence and multiple glacial refuges have been found for several species [46, 49–51]. In the case of the populations of *Ceratozamia* in the lowlands (*C. brevifrons*), we found that the species took refuge in the foothills of the Sierra de Chiconquiaco during glacial cycles.

There is evidence to suggest that *Ceratozamia* species found refuges in southern Mexico due to the climate changes of the Pleistocene, where the greatest number of basal clades and diversity of the genus are currently concentrated [47]. Though, there are no Pleistocene refuges in the areas north of the Trans-Mexican Volcanic Belt [47]. What according to shows that *Ceratozamia* expanded from southern Mexico, so the species north of the Trans-Mexican Volcanic Belt are of recent speciation [10,47,52]. The stage of climatic changes that influenced the distributions of the cycad species in this study, whose data come from the late Pleistocene, coincide with the origin of some species of the “Mexicana” clade, including *Ceratozamia tenuis* whose origin dates to the middle Pleistocene (1.4 my) [10]. Therefore, these reductions in the distribution of the species that were recently separated at that time, contributed to the isolation of their populations avoiding subsequent interactions, which continues to the present since the three species are in different ecoregions.

The diversification of *Ceratozamia* clades in the Pleistocene occurred in the mountainous region of the Sierra Madre Oriental and southern Mexico and even Central America [10]. Furthermore, with the information generated through the ecological niche models, we inferred a reduction in habitat suitability in the past of at least these three *Ceratozamia* species studied concerning the present. According to the Pleistocene refuge hypothesis, these species in the postglacial periods had an increase in their distribution up to the present time and are developing in secondary contact areas [53]. However, these cycad species did not have moments of later encounter because their current distributions are separated by notable orographic barriers, being a good example of allopatric speciation.

At present, there are remnants of populations concentrated in ravines with difficult access where agriculture and livestock activities are not profitable in Veracruz. Even under these conditions, the reduction of its population is continuous although on a smaller scale due to livestock activities (e.g., goats and cows) and seasonal crops on a smaller scale. We recognized according to the evidence that what remains today are relict populations of species whose lineage has developed through millions of years, enduring different climatic events of different intensities [27], which has placed them in different international and Mexican protection schemes [5, 54].

In addition to the above, we showed that future modelling indicates that their surface will be reduced to such a degree that they practically disappear, especially in the cases of *Ceratozamia morettii* and *C. tenuis*. A previous study has shown that in the case of *Ceratozamia miqueliana*, future modeling does not cause any change in the suitability of its niche (Carvajal-Hernández, unpubl.), however, it is a lowland species subjected to temperatures that exceed 35° C in the summer season [25]. This contrasts with what is shown in the models of these species (*C. morettii* and *C. tenuis*), which are found in mountainous regions with temperate temperatures, but it matches the future forecast of *Ceratozamia brevifrons* whose model indicates an increase in its distribution.

There are models that indicate that in Mexico temperate ecosystems (coniferous forest, *Quercus* forest and cloud forest) will suffer a reduction due to climate change, while tropical ecosystems will have an increase in their distribution [55]. This coincides with our results, since the species that are estimated to have a drastic reduction are precisely those that inhabit temperate forests, while *C. brevifrons* currently lives in conditions of higher temperatures and is subject to greater droughts, it will have an increase in its distribution. A similar case was registered with several *Pinus* species in Mexico with neartic affinity, emphasizing that for *Pinus oocarpa* a decrease in its distribution of almost 97% is predicted, in contrast to *Fuchsia microphylla*, a topical species whose distribution would increase considerably in the future [56]. This situation has been described with other temperate species, attributing changes in the distribution associated with the loss of humidity, assuming then that according to their physiological tolerances, the species will be affected in the short term by a reduction in humidity rather than by a temperature increase [57].

Climate change models indicate that tropical mountains will be the systems most affected due to the loss of species [58], and therefore its contained biodiversity, which is higher compared to lowlands in biodiverse tropical zones [59]. It is estimated that the mountains will become warmer and drier, with a trend towards greater drought in the lower mountainous areas which will cause changes in the distribution of plant species, some will benefit but others will be harmed in various aspects [60,61]. In some mountains of the world, the ranges of distribution of plant species will be shortened as a result of climatic changes, droughts in low areas and extreme snowfall in high areas [60]. According to the habitat suitability models, the cycad species studied here are likely to be affected by apparent droughts, since at present both *Ceratozamia tenuis* and *C*. *morettii* are found in sites with high humidity and temperate temperatures. However, the future geographic distribution of *C*. *brevifrons* seems to be affected to a lesser extent, perhaps because the current distribution of the species corresponds to dry climates, which may represent an adaptive advantage in the future.

## Conclusion

We consider that the three species of cycads are threatened, due to their limited area of distribution that makes them microendemic, in addition to the anthropogenic pressures on their habitat. Climate models of the past indicate an isolation of their populations, same that continues to the present. Through future climate models, a drastic reduction in the suitability of the habitat of *C. morettii* and *C. tenuis* that currently inhabit temperate mountainous regions is expected, a situation that increases their current vulnerability. On the contrary, an increase in distribution is estimated of *C. brevifrons* that now lives in conditions of higher temperature and drought. These models show the trajectory of the species through time, observing that they have endured different climatic changes that affect their distribution. However, current anthropogenic pressures, added to future models, indicate that two of these species will be susceptible to disappearing in a short time, both due to climatic trends that will affect their habitat and due to anthropogenic pressures, that are expected to continue in their distribution area. However, it is interesting to note that future climate change will affect the distribution of species differently, such is the case of *C. brevifrons*, whose current habitat conditions will apparently be maintained and will even increase favoring its possible distribution. This raises different scenarios for biodiversity in general.

Cycads represent interesting species from a biogeographic perspective due to their restricted distribution, but for that reason, they also represent a challenge for their conservation. There is no doubt that the understanding and recognition of the factors and patterns that originate biodiversity on temporal and spatial scales will help the conservation strategies of ecosystems and species to be successful.

Unfortunately, how habitats and species in general can respond to climate change is not yet fully known, so there is still an urgent and challenging task to fill these information gaps. Therefore, to guide conservation efforts efficiently and in future ecological studies, multiple perspectives and approaches are necessary to be integrated, as well as to know the complete history of the groups to be studied as in this study.

## Acknowledgments

We thank Mohamed bin Zayed Species Conservation Fund and the Centro de Investigaciones Tropicales (CITRO), Universidad Veracruzana for their support for conducting fieldwork. We also thank Biologist Hector Hernández Andrade for his permission to carry out fieldwork on his property and to Merbin Tornero Conde for his support in the field.

